# Sensory and palatability coding of taste stimuli in cortex involves dynamic and asymmetric cortico-amygdalar interactions

**DOI:** 10.1101/2025.07.01.662567

**Authors:** A. Mahmood, J.R. Steindler, D.B. Katz

## Abstract

Gustatory cortical (GC) and basolateral amygdalar (BLA) taste responses consist of an inter-regionally coherent 3-part state sequence. This coherence suggests that reciprocal BLA-GC connectivity is important for taste processing, but it remains unknown: 1) whether BLA-GC coherence actually reflects a reciprocal “conversation” (as opposed to one region simply driving the other); and 2) whether such a “conversation” has anything to do with the taste processing observed within GC response dynamics. Here, we address these questions using network and single-neuron analysis of simultaneously-recorded GC and BLA taste responses in awake rats. We find asymmetric, reciprocal µ-frequency influences that reflect taste processing dynamics: BLA→GC influence dominates between 300 and 1000msec (the epoch in which BLA codes palatability); afterward, when GC responses become palatability-related and GC has been shown to release a behavior-relevant signal, the direction of influence reverses, becoming GC→BLA. Follow-up analyses demonstrate that this “turn-taking” exists alongside effectively synchronous amygdala-cortical coupling—the two regions functioning as a unified structure. Finally, to assess the implications of these interactions for single-neuron responses, we tested the response properties of GC neurons categorized by their inferred connectivity with BLA: GC neurons influenced by BLA produce stronger taste-specific and palatability-related responses than other GC neurons, and the strongest taste encoding is specifically found in GC neurons that both influence and receive influence from BLA—those most deeply embedded in the reciprocal circuit. These results, consistent with findings in multiple systems, support the novel conclusion that taste processing and decision-making is a function of the amygdala-cortical loop.

**New & Noteworthy:** Conventionally, taste circuitry is considered feedforward, travelling up from the brainstem, with each additional node containing more sophisticated information. We challenge this convention by demonstrating that amygdala and cortex instead influence each other bidirectionally, yet in a direction-specific and asymmetric manner. These influences appear to drive different parts of the taste response, with distinct patterns for decision-making and behavioral output; furthermore, involvement in cortex-amygdala functional connectivity determines the strength of encoding in cortical single neurons.

## Introduction

Stimulus-evoked activity in gustatory cortical (GC) neurons transitions through a sequence of population states, with each successive state containing more highly processed information (1,2). Across the 1-1.5 sec following taste delivery to the tongue, these states reflect, in turn, the fact that the taste is on the tongue, then taste identification, and finally the decision to either consume or reject the taste on the basis of its perceived palatability (3,4). Such ensemble neural dynamics are predicted to arise in systems with dense recurrent connectivity (5,6), which has been shown, in studies on visual processing, to generate dynamic neural responses observed across multiple regions, as inter-region influence cascades in forward and reverse directions across the visual stream (7–9).

Another expected consequence of reciprocal inter-region connectivity is coupling of activity across regions. In line with this expectation, recent work has shown both correlations in the latencies of the above-described firing-rate transitions and significant cross-coherence between GC and Basolateral Amygdalar (BLA) taste responses (10), both which have strong reciprocal connections (11–15). these findings dovetail nicely with work demonstrating cross-coherence between reciprocally connected structures in other sensory systems (16,17), together suggesting that the reciprocal connectivity between BLA and GC may be central to processing of the taste itself.

Current literature stops well short of testing that hypothesis, however. Previous studies on inter-regional interactions in taste processing have largely focused on the impact of perturbing one region on processing in the other (4,18–22), and while a few studies have investigated multi-region recordings, they have done so using cross-coherence measures (10), usually collapsed across time (19,23). Since these techniques are agnostic to which connections cause “coupling,” they cannot distinguish unidirectional from bidirectional influences. That is, it is unknown whether the evolution of the neural processing of tastes reflects moment-to-moment changes in which region’s activity is “influencing” that in the other.

The current study answers these questions by bringing advanced analyses to bear on simultaneously-recorded BLA and GC taste-evoked activity (in a paradigm that avoids confounds related to GC’s involvement in interoception, see Methods). We show that the BLA-GC interaction is a conversation in which the two regions take turns influencing one another, and that this asymmetry of influence is epoch-specific: BLA primarily influences GC during the 300-1000ms post-stimulus period, which is the epoch during which BLA responses are palatability-related (10,23,24), and just before GC responses become so (2,3); GC influence on BLA then peaks briefly at approximately the time of the transition into the next epoch—the point at which GC has been shown to emit behavior-related signals (25). We perform follow-up analyses to show that this “turn taking” asymmetry exists within the framework of nearly synchronous, coordinated dynamics—as expected for a system in which function relies on reciprocal interactions.

We then move on to characterizing the functional properties of single neurons underlying these interactions, testing the hypothesis that embeddedness in the amygdala-cortical loop determines involvement in taste processing. We show that roughly half of our GC single-neuron sample can be inferred to either receive input from or send output to BLA. These overlapping functional “input” and “output” populations are both primarily made up of putative pyramidal neurons, and yet are distinct with regard to their roles in taste processing: GC neurons receiving influence from BLA show significantly higher levels of both taste specificity and palatability-relatedness in their responses than those that only influence BLA; furthermore, GC neurons that both influence and receive influence from BLA produce the strongest taste-related responses.

Collectively, these results shed light onto the forces underlying taste-related processing by demonstrating that dynamic recurrent interactions between BLA and GC play a role in the emergence of GC neural codes related to the generation of the behavioral response, and that the degree to which a GC neuron is embedded in the reciprocal circuit largely determines its involvement in taste processing. These results highlight the importance of inter-region interactions in understanding perceptual function.

## Results

### BLA-GC interaction dynamics are asymmetric but reciprocal

Insight into the involvement of reciprocal amygdala-cortical coupling in taste processing can only be gained through an examination of the dynamics of directional influence—of the moment-to-moment degree to which activity in each area impacts activity in the other—and of the degree to which this interplay reflects the known taste processing dynamics. We began this examination by applying Spectral Granger Causality (GrCa) to local field potentials (LFPs) simultaneously recorded from BLA and GC. Rather than simply assessing similarity between timeseries (as cross-coherence analyses do), GrCa allows us to disentangle directional dependencies by making use of time-lagged regressions: if future GC activity is better predicted using the histories of both GC and BLA, compared to just using the history of GC, then BLA can be thought to “causally influence” the future of GC (26–30). Significance of influence in GrCa can then be assessed for each time and frequency bin, and thus we can test whether GC and BLA are mutually influencing one another (and whether those directional influences have different time-courses). We note that GrCa infers functional connectivity between regions, and neither has information regarding, nor makes claims about, the physical connectivity between the sources of its data; hence, we use the term “influence” here to describe only a functional connection without claiming any specific properties of a neural projection (excitatory or inhibitory).

**Fig. 1** shows the basic (collapsed across time) output of this analysis brought to bear on simultaneously recorded GC and BLA LFPs. The influence of each region on the other is almost entirely contained within the 1-60Hz range (**Fig. 1A**; note that the low-pass filter used when collecting the LFPs allows for the detection of influence at much higher frequencies, had they existed, see Methods). More to the point of our investigation, the GC→BLA influence proved quite distinct from the BLA→GC influence within that range (**Fig. 1B**): GC→BLA influence, while above zero across this entire range of frequencies, dominates in the 20 to 60Hz range (β and γ frequencies; see Ref# 31); BLA→GC influence, meanwhile, is essentially non-existent (and non-significant) across most of the β/γ range, but is the dominant direction of influence in the θ and µ frequency ranges (2-12Hz), the latter of which is sometimes referred to as α (31,32). This latter frequency range is known to be both particularly prominent in GC evoked activity and modulated in a manner related to the dynamics of GC taste responses (31–33). In summary, BLA influences GC and GC influences BLA during taste processing, and these two influences are distinct from one another.

**Fig. 1).**
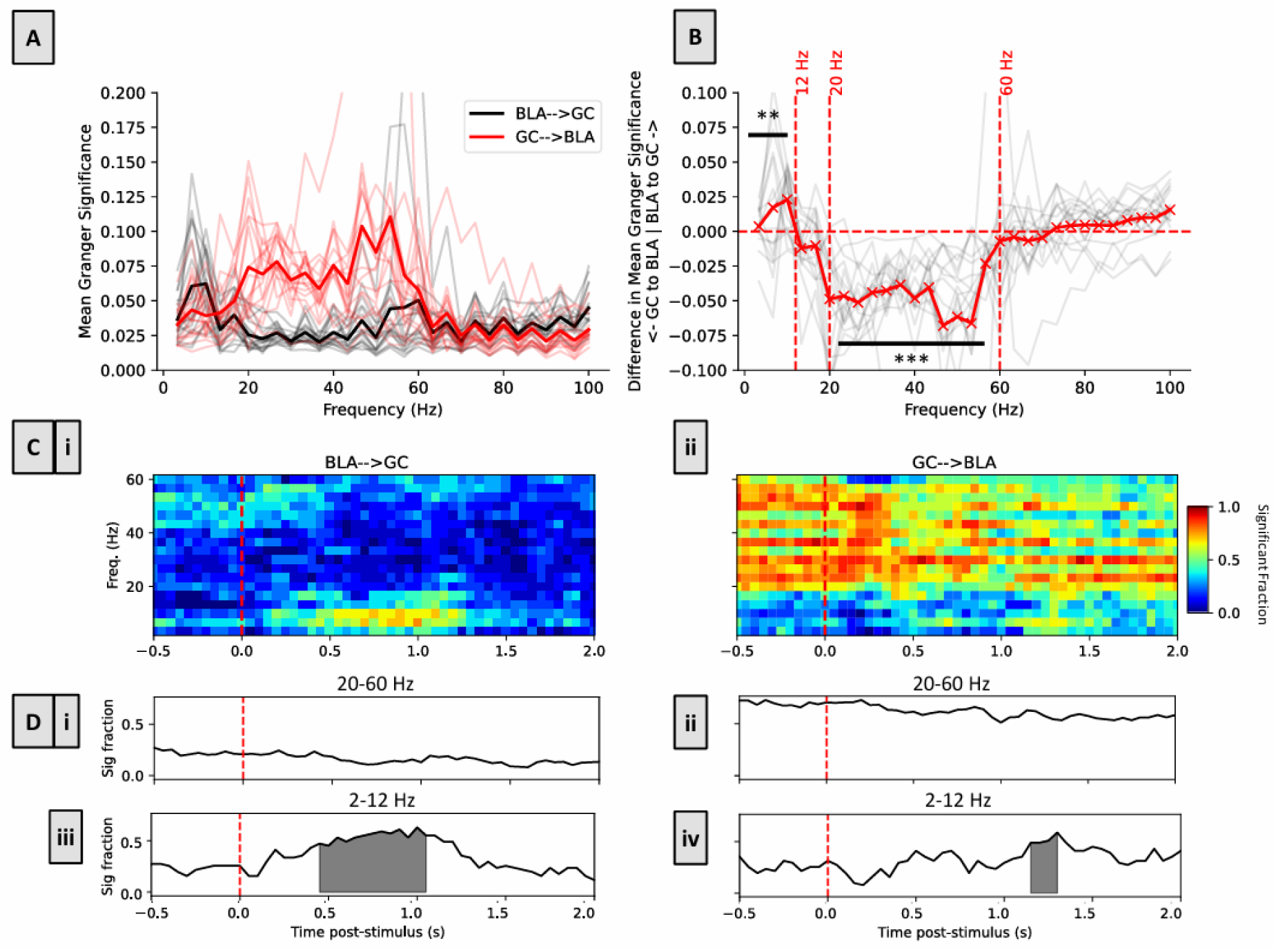
GC-BLA interaction is rich with processing-related asymmetries and dynamics. **A)** Mean (bold lines) and individual sessions’ (thin lines) significance of Spectral Granger Causality for 0-2000ms post-stimulus activity across datasets for BLA→GC (black) and GC→BLA (red) influences. **B)** Directional differences in mean (bold red line) and individual subjects’ (thin black lines) influence across frequency bands for data shown in A). Above-zero differences mean stronger BLA→GC influence, and below-zero differences mean stronger GC→BLA influence. BLA→GC is stronger below 12Hz, and GC→BLA dominates between 20 and 60Hz; ** = p<0.05, *** = p<0.005. **C)** The data from Fig. 1A unpacked into significant time-frequency bins for **i**) BLA→GC and **ii**) GC→BLA influence. Brighter color indicates that a larger fraction of the recording sessions displayed significant Granger Causality at that time-frequency bin (heatmap legend to right). In both panels the difference between higher and lower frequency bands can be seen, and across panels the asymmetry in influence can be seen; furthermore, dynamics of influence are apparent in the lower frequency band. **D)** The data shown in C collapsed across frequencies within each identified bandwidth (frequency band limits are given above each panel). Dynamics of influence are restricted to the <12Hz bandwidth; shaded regions indicate periods of strongest significance determined using cluster-permutation testing. See text and Methods for more details.

We note that it is unlikely that this relatively clean separation in directional frequencies is an artifact of the methodology, as GrCa can appropriately recognize influences in identical or overlapping frequency bands, as has been demonstrated before (26,34–36). Furthermore, the distinctness between the frequency bands of the BLA→GC and GC→BLA influences aligns with previous work showing that pharmacological inhibition of taste thalamus (VPMpc) reduces GC LFP power in the 20-60Hz band, but increases it in the 2-12Hz band (19)—a finding that corroborates the idea of the 2-12Hz and 20-60Hz bands as being distinct communication “channels”.

The distinction between lower and higher frequency ranges is also easily seen when direction-specific influence is examined across time (**Fig. 1C**; compare the upper two thirds to the lower third in each panel), as is the difference between GC→BLA and BLA→GC influence (compare the left and right panels). This presentation further suggests that, in general, the β/γ influence may be slightly higher spontaneously than during taste processing. But most centrally with regard to our hypothesis, the differences between BLA→GC (**Fig. 1Ci**) and GC→BLA (**Fig. 1Cii**) influences within the θ/µ frequency range can be seen to change repeatedly across taste processing time, and this “turn-taking” reflects the oft-described GC dynamics.

To rigorously assess the significance of this latter phenomenon, we plotted the average proportion of sessions in which power in each frequency-time bin is significantly above zero (the “significant fraction”) in each direction of influence, collapsed across frequencies within each band. This analysis reveals near-tonic influences in the higher frequency band (**Fig. 1Di** and **ii**; the average GC→BLA magnitude of influence is of course higher than that of BLA→GC influence), although there is a slight trend toward a reduction of influence following taste delivery.

The µ/α band results, meanwhile, do in fact reflect the known GC taste-response dynamics (**Fig. 1Diii** and **iv**). In each panel, the shaded regions represent the result of cluster-based permutation testing, identifying the longest continuous periods in which this result is significantly higher than that of randomized data (note that there are no such periods for β/γ influence; **Fig. 1Di/ii**). With regard to BLA→GC influence in the lower frequency band (**Fig. 1Diii**), this period is a good match to the GC identity epoch (approx. 300-1000ms post-stimulus; see **Fig. 3A**). Given that BLA responses are palatability-related during this period (24), and that GC responses become palatability-related only at the end of this period, this result also accords well with classic studies on the role of amygdala in stimulus valuation, which have led both human and rodent researchers to theorize that amygdala “passes” emotional valence to cortical regions (4,18,37,38) see also Discussion).

GC→BLA influence in this lower frequency range, meanwhile (**Fig. 1Div**), peaks only briefly (note the thin string of yellow bins between 1.0 and 1.5sec in the bottom rows of **Fig. 1Cii**), with that peak coming immediately following the BLA→GC push. This influence “turn taking” further corroborates the above interpretation, suggesting that GC, after being influenced by BLA, emits decision-related activity capable of influencing behavior, consistent with work demonstrating that the transition into palatability-related firing in GC (which typically occurs around this time) is causally linked to the driving of behavior (3,25). This result establishes bi-directional interactions between BLA and GC during taste processing, with dynamic asymmetries of amygdala-cortical coupling that directly reflect what is known of GC taste processing.

### Asymmetric BLA-GC interaction dynamics are nonetheless synchronized

Again, the fact that this amygdala-cortical “conversation” is asymmetric was predicted (see Discussion section “The division of function(s) between BLA-GC directional influences”). And while it might seem at first blush that they contradict the previously-described coherence of activity in the two structures (10), further analysis revealed that even these asymmetries of influence reflect the fact that the two regions process tastes according to the same “clock”—that changes in influence occur simultaneously. This can easily be seen when Principal Component Regression is performed on each time-series shown in **Fig. 1C**. In **Fig. 2A**, the trajectories for the data in the two Fig. 1C panels obtained from this regression are shown overlain on one another. The dynamics of the two time-series, which are represented as changes in the trajectories’ directions through PCA space, are clearly very similar.

**Fig. 2).**
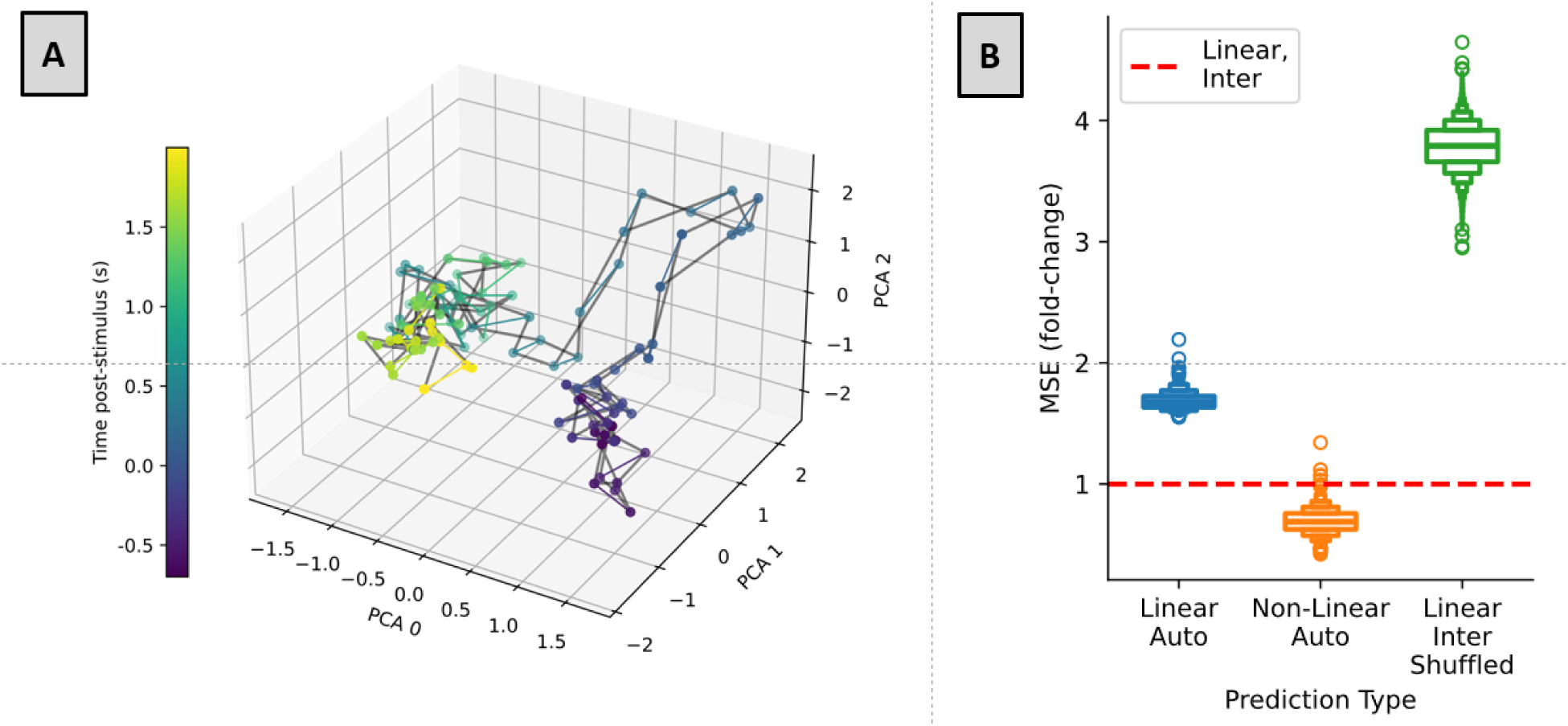
**Dynamics of BLA**→**GC and GC**→**BLA influences are time-locked. A)** Aligned principal components of spectra shown in Fig. 1C, indicating that the dynamics of BLA→GC and GC→BLA influence are strongly coordinated, in that the trajectories change direction together through time. Elapsing post-stimulus time is shown with changing colors (legend to left). **B)** Quantification of alignment shown in A). Prediction of one time-series (GC→BLA) from the other (BLA→GC) using linear regression (Linear, Inter) is better than both shuffled linear regression (Linear, Inter, Shuffled), and linear autoregression (Linear, Auto) of timeseries, and almost as good as non-linear autoregression (Non-linear, Auto).

**Fig 3.**
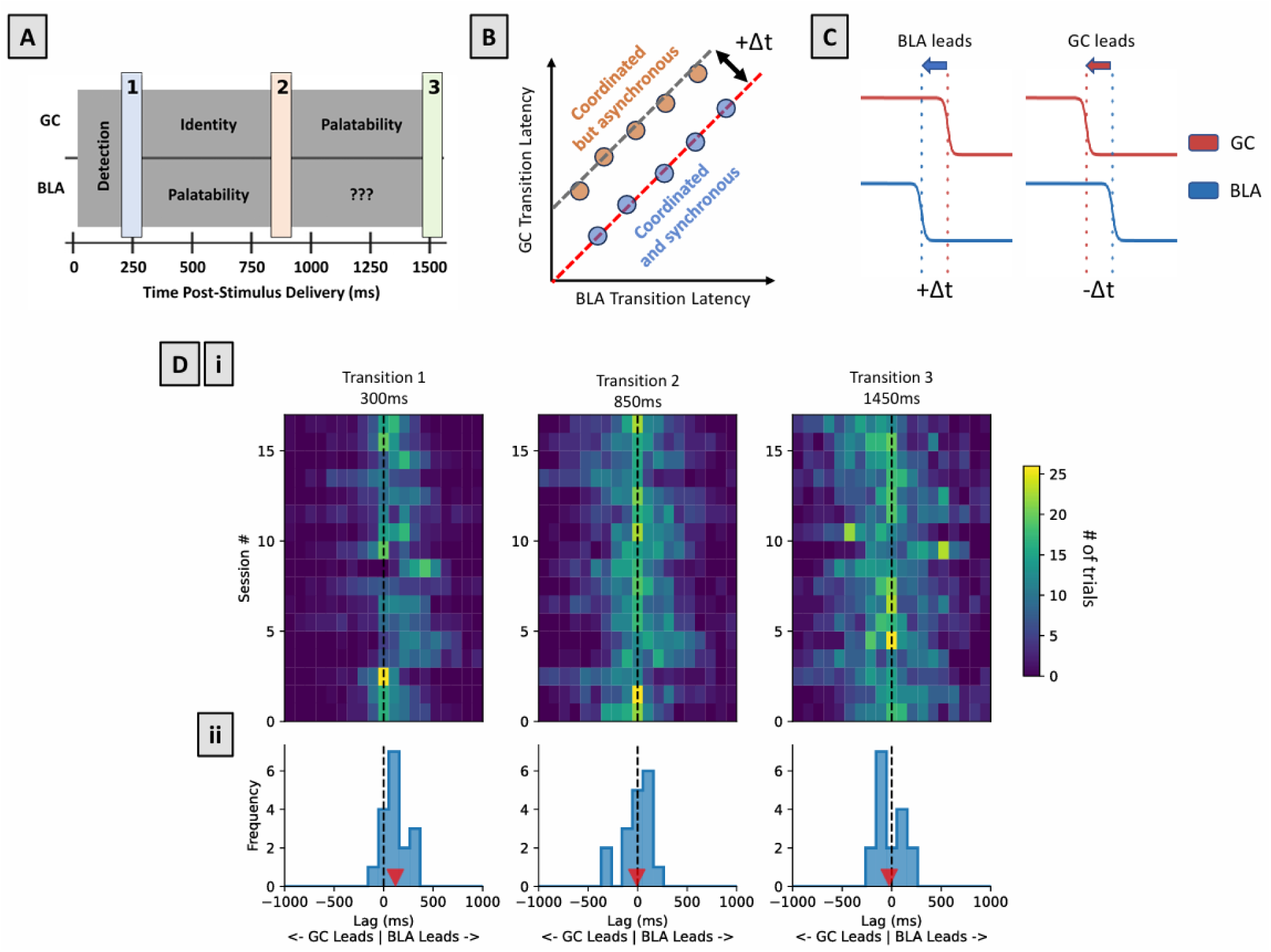
BLA initially leads GC in state transition but both regions subsequently show zero-lag coordination. **A)** Summary schematic of the timing and “coding content” previously demonstrated to characterize state sequences in GC and BLA ensemble spiking activity. Overlain are the numbers used in this study to label these transitions. **B)** State transitions can be strongly coordinated but may still show significant lags. **C)** A schematic illustrating how transition lags between GC and BLA were calculated and how we define the sign of the lags. **Di)** Distributions of transition lags, presented as heat maps (legend at right shows the # of trials with a specific transition lag) for 18 sessions (y-axis) across 6 animals for each of the 3 transitions. **Dii)** Summary distributions of lags for each transition, with red arrows indicating the grand-average lag. The significance of the difference between that grand-average lag and 0 was evaluated using bootstrapped 95% confidence intervals of the distribution mean (see text and Methods).

We quantified this similarity by assessing the mean-squared error (MSE) of attempts to linearly predict one time-series from the other (**Fig. 2B**, red dashed line) compared to that produced by the same linear regression of the two time-series after shuffling them temporally (linear, inter, shuffled), and also by linear and non-linear (i.e., using multi-layered artificial neural network) auto-regression (i.e., predictions of each time series from itself; see Methods), which give us a lower bound for prediction error. The results of these analyses make it clear that the linear prediction of the BLA→GC time-series from the GC→BLA time-series (in blue) was almost as accurate as the non-linear auto-regression (orange) and better than the linear auto-regression (green; this was not surprising given the nonlinearities in the BLA-GC interaction dynamics). Simply put, the inter-series prediction nears the limit of prediction accuracy, a fact that serves as novel evidence for the suggestion that BLA and GC form, upon receiving taste input, a single distributed dyad with time-locked, coupled dynamics— albeit dynamics within which GC→BLA and BLA→GC influences are asymmetric.

As a convergent measure, we tested the specific timing of the coordination between the ensemble firing-rate state changes (**Fig. 3A**) that have been repeatedly identified in GC and BLA (10,24). While our previous results show the latencies of GC and BLA state changes to be correlated, this leaves open the question of whether there are significant lags (or offsets) between ensemble firing-rate transitions in either region (see **Fig. 3B**), which would suggest a general direction of “influence flow.” We therefore directly assessed these lags (see **Fig. 3C** for a schematic of the analysis).

The between-region lags for each individual session (**Fig. 3Di**, presented as a heat map) appear to be highly reliable across the 18 sessions (from 6 rats), appearances that are confirmed by the tightness of the across-session distributions of mean lags (**Fig 3Dii**). Analysis of these latter distributions reveals that, for the earliest transition, the sudden transition in BLA ensemble firing rates tends to lead that in GC (permutation test, 95%CI=[57, 183], p<0.001), but fail to show significant between-region deviation from simultaneity (zero-lag) of later transitions (Transition 2: 95%CI=[-74.5, 51.5], p=0.78; Transition 3: 95%CI=[-86, 39.5], p=0.43). The distributions of lags for transitions 2 and 3 were both significantly different from that for transition 1 (1 vs. 2: p=0.001, 1 vs. 3: p=0.015, Wilcoxon signed-rank test), but not from one another (Wilcoxon signed-rank test, p=0.747). This finding agrees with previous results which showed that the first transition is not significantly correlated between BLA and GC, but the 2^nd^ and 3^rd^ transitions are (10).

These results support the hypothesis that BLA and GC are not a joint processing network “at rest,” but that they coalesce into a dyad—not just working together, but becoming essentially a single processing object—after producing uncoordinated initial transients in their taste responses (see Discussion). The lag present in the first transition suggests that BLA and GC taste responses are initially driven independently (and provides proof that the analysis is capable of detecting even small reliable lags), although there are other possible explanations. Regardless, this initially uncoupled activation gives way to the two regions cycling through the later transitions of their taste responses as a single processing unit, such that the relatively long axons connecting the two regions do not induce detectable delays in the firing-rate transitions that characterize taste processing (see Discussion).

### Modeling of single-neuron functional connectivity reveals overlapping groups of GC neurons differentiated on the basis of whether they influence or are influenced by BLA neuron activity

As noted above, the finding that different patterns of influence appear to “live” in different frequency bands is not without precedent—bottom-up and top-down influences in visual pathways have also been suggested to segregate into separate frequency bands (35,36,39). Meanwhile, the fact that BLA→GC influence is focused in the 2-12Hz range provides further motivation for our central hypothesis that the involvement of GC neurons in taste processing (which can be quantified in terms of the magnitude of GC responses’ taste-specificity and palatability-relatedness) might be a function of their degree of coupling with BLA, given that previous studies have shown that GC taste-processing dynamics are also prominently visible in GC µ activity (19,31–33).

To set up the test of our central hypothesis, we trained Poisson Generalized Linear Models (pGLM, see Methods for important differences between pGLM and the more commonly-used General Linear Model) to infer connectivity and influence between individual GC-BLA neuron pairs—an aim that cannot be achieved with GrCa (see Methods), and that allows us to categorize individual GC neurons as putatively influencing and/or being influenced by activity in BLA (and vice-versa). We fit pGLMs to each single neuron in our dataset, using spike trains from all other simultaneously recorded neurons as input (40–42).

An advantage of this analytic approach to inferring functional connectivity is that, while it cannot capture the dynamics of influence, it is inherently more conservative than correlational methods such as spike-train cross-coherence (43). Standard correlational approaches are significantly more prone to false positives due to the shared latent dynamics across neurons within regions and across regions (41). In pGLMs stimulus input and the history of spiking in the current neuron “compete” against one another, and also against the recent spiking of other neurons (**Fig. 4A**), for significance during pGLM inference. The evoked activity of a neuron can potentially be explained as a function of any of these variables, and only explanatory value above and beyond what is predicted by all other variables is deemed evidence of a functional inter-regional influence—influence that can be attributed to other sources is essentially “partialled out.”

**Fig. 4).**
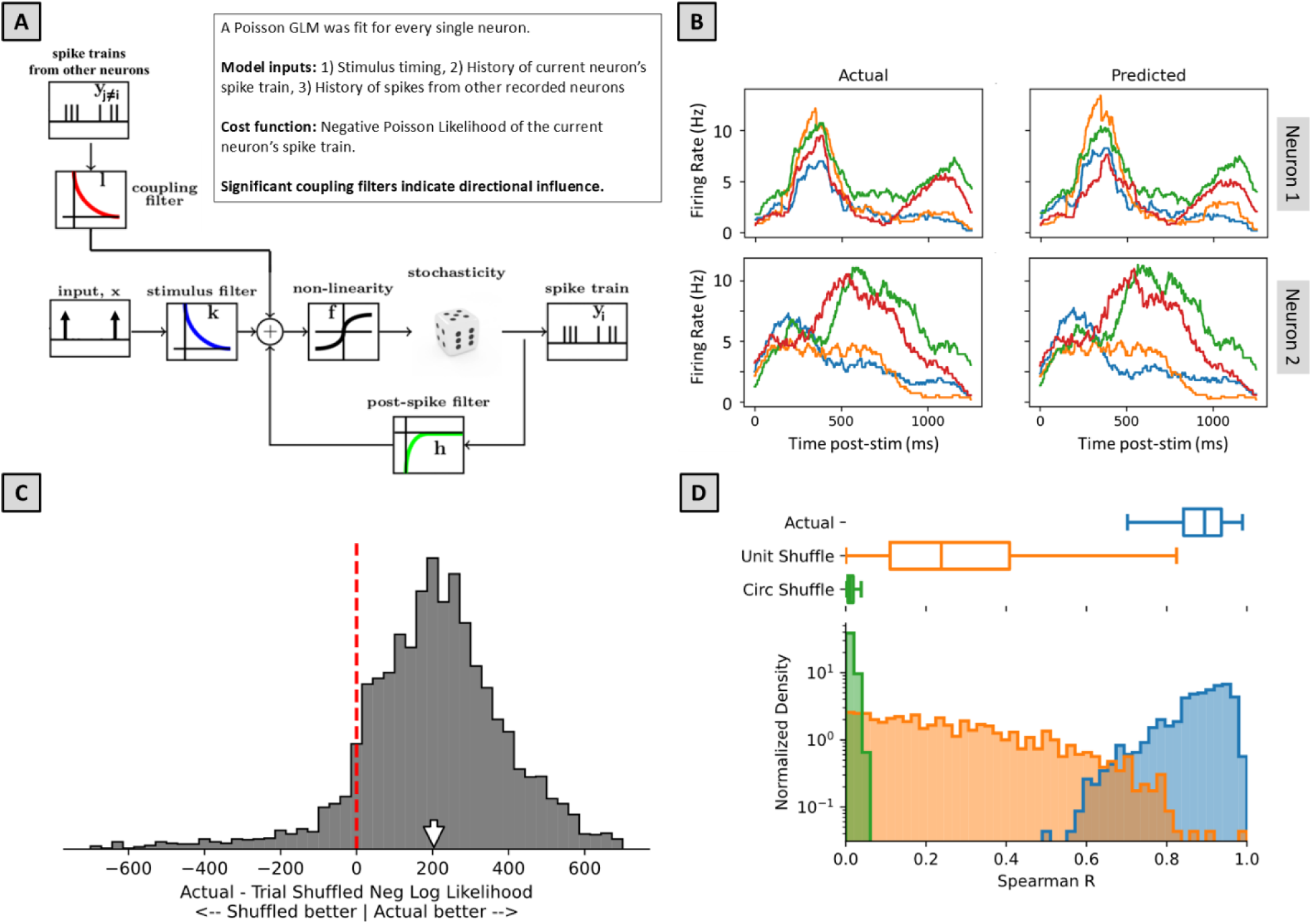
Poisson Generalized Linear Modelling of single-neuron interactions between BLA and GC provides good fits to the data. **A)** pGLM structure. Every neuron model depends on: 1) stimulus timing, 2) the neuron’s spiking history, and 3) input from (or spiking history of) all other simultaneously recorded neurons. **B)** Actual and Predicted PSTHs of two representative units across 4 delivered tastants (different colors), showing that fitted models (right column) accurately recapitulate single-neuron activity (left column). **C)** The vast majority of models built on real data showed higher cross-validated likelihood than models trained on shuffled data (i.e., the bulk of the distribution is to the right of the vertical red dashed line at equal predictability). This confirms that the models are learning single-trial relevant features. The white arrow indicates the median of the distribution. **D)** The distributions of correlations (Spearman’s Rho) between: 1) actual and predicted PSTHs (blue); 2) unit-shuffled data and prediction (orange); and 3) temporally scrambled data and prediction (Circ shuffle: green).

In total, we fit models to 151 and 160 well-isolated GC and BLA single neurons, respectively (mean±SD per session: 9.4±4.14, GC; 10.0±4.61, BLA). Before using model output to test any hypotheses about taste processing, we tested the quality of the fit to the data provided by the models. **Fig. 4B**, 2 representative examples of actual PSTHs (left) and PSTHs predicted by the pGLMs (right), visually illustrates this fit. The apparent similarity between the two is quantified in **Fig. 4C**, which shows that the model consistently produces significantly better matches to the data than models fit on spike-trains shuffled between trials (Wilcoxon signed-rank test, p < 0.001), a control that eliminates only the within-trial coordination of spiking between simultaneously recorded neurons, preserving the average statistics of the spike-trains used. This result highlights the fact that, despite significant trial-to-trial variability in spiking, the pGLM learns within-trial relationships that are meaningful.

As an additional test of the pGLM fits, we also compared correlations between actual PSTHs and: 1) trial-matched predictions; 2) cross-unit predictions; and 3) predictions on circularly shuffled data (**Fig. 4D**, see Methods for details). We found, as expected, that trial-matched predictions (blue histogram and summary box plot) perform best. Control correlations with unit-shuffled/cross-unit predictions (in orange) were significantly worse (Wilcoxon signed-rank test, p<0.001) but also significantly above zero, probably because many GC neuron pairs exhibit broadly similar neural dynamics. Worst, and essentially zero, were correlations with circularly shuffled (temporally garbled) data (in green). Together, these tests allow us to conclude that the pGLM generates reasonable models of the mechanics (functional circuitry) underlying the observed activity.

These results confirmed that we can confidently use the model to evaluate whether the activity of one simultaneously recorded neuron influenced that of another (i.e., we could identify significant coupling filters). In what follows, we for simplicity’s sake refer to neurons with responses that are concluded to exert influence on activity in the other region as “Output” neurons, and neurons with responses that are influenced by activity in the other region as “Input” neurons; note that in neither case are we proposing to have identified direct anatomical connections—the terms refer purely to functional connectivity.

**Fig. 5A** shows the distribution of neurons so identified: clearly, our BLA sample contains more Output neurons (40.0%) than Input neurons (6.3%), with an additional subset (9.4%) categorized as both; the Output and Input populations in GC were more balanced (17.2% and 22.5%, respectively), with an additional 11.9% characterized as doing both. Another subset of neurons in GC could not be described as either influencing or being influenced by activity in BLA (and vice-versa). No claim is made about whether these neurons might output to or take input from any structures from which we didn’t record, and therefore no further analysis is done on the (probably highly diverse set of) neurons that were categorized as neither Input nor Output.

**Fig. 5).**
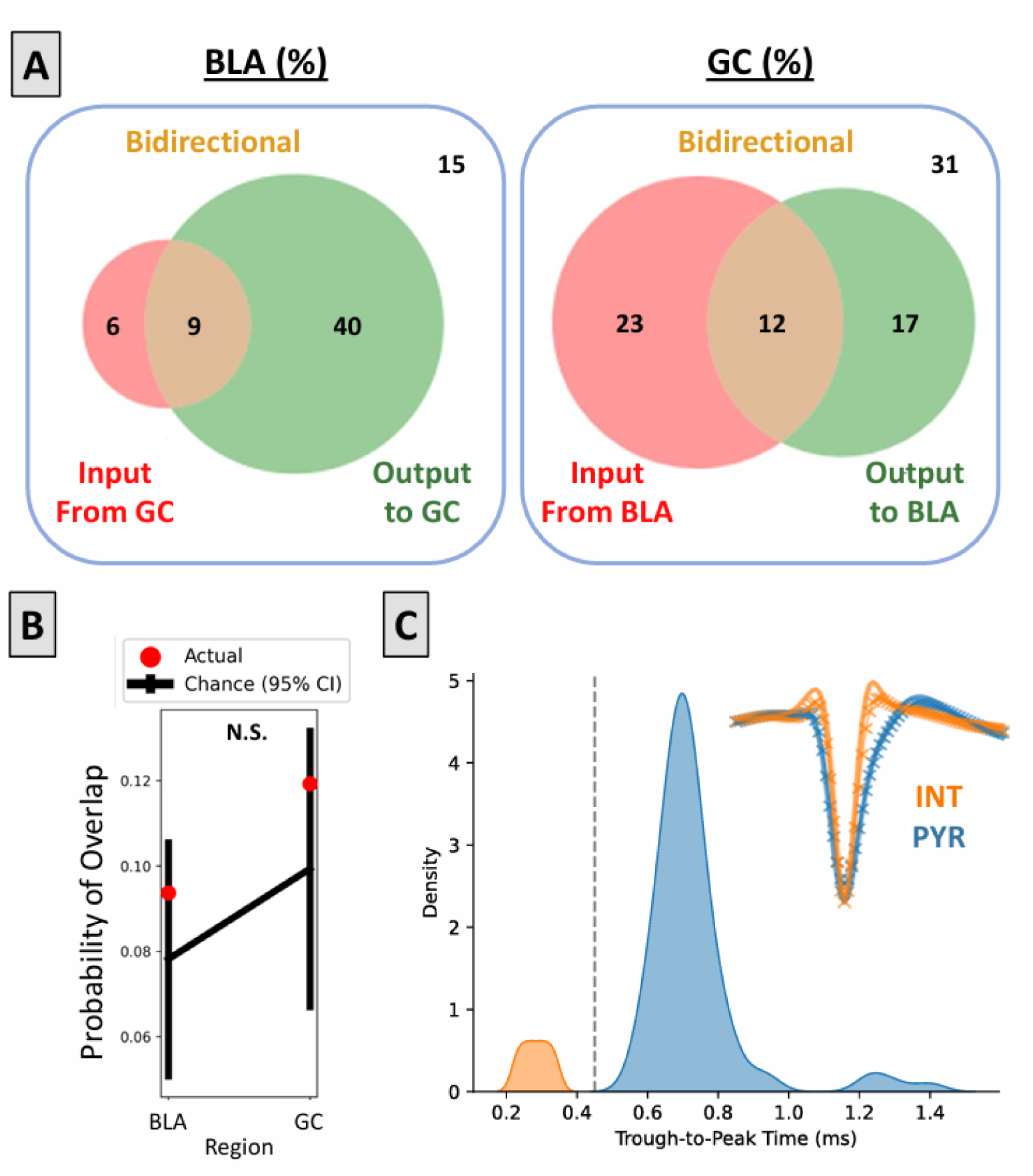
pGLM brought to bear on amygdala-cortical ensembles identifies functionally (but not structurally) distinct subpopulations of neurons. **A)** Inference of Input and Output population in the recorded BLA-GC populations (number outside colored circles indicates percentage of neurons with no identified between-region connections). **B)** The amount of overlap between Input and Output populations is not significantly different from that expected by chance in either brain region. **C)** Inferred neuron-types based on trough-to-peak times of waveforms (Inset: solid line = GC, crosses = BLA). Most neurons in the dataset (expectedly) are putative pyramidal neurons, which means that the vast majority (and similar percentage) of each functional neural subtype as shown in panel A is pyramidal.

Given the relatively small overlap between sets of functional Input and Output neurons shown in **Fig. 5A**, it was reasonable to ask whether the two sets were distinct—that is, whether a neuron (or neuron type) was specifically likely to be either an Input or Output neuron, but not both; as has been suggested for projection neurons in the visual stream (17,44). Two separate analyses provided evidence against this possibility. First, we performed a direct, Bayesian assessment of the overlap between Output and Input sub-groups in both regions, comparing the data to a null distribution in which labels for connection types were assigned randomly so that we could ask whether, given the likelihood of a neuron being categorized as Input or Output, the conditional probability (i.e., the likelihood of it being categorized as Input given that it was categorized as Output, and vice-versa) was lower or higher than chance. **Fig. 5B** shows that the overlap we observe in the real data falls squarely within the chance probability of overlap for this null-distribution for both regions, therefore providing no evidence that Output and Input populations should be thought of as segregated groups.

We also assessed the likelihood that the separation of neurons into Output and Input groups aligns with anatomically-defined neuron categories. Specifically, we asked whether putative pyramidal cells, identified on the basis of spike width, were more likely to be Output neurons than Input neurons. **Fig. 5C** demonstrates that our sample of isolated single neurons broke, as expected, neatly into two groups on this basis, with a 0.45ms trough-to-peak time providing a clean cutoff between fast-spiking putative interneurons (Int) and slower-spiking putative pyramidal neurons (Pyr; see (4,45–47). Slightly under 10% of our sample fell into the Int category, a number that is not wildly inconsistent with expectation based on an extensive previous literature using chronic extracellular electrophysiology. Given the small fraction of interneurons, it can be concluded that putative pyramidal neurons are approximately equally likely of being Input and Output neurons. In conclusion, we could find no evidence suggesting that Output and Input neurons are distinct anatomical types (at least at the level of gross subdivision).

### Taste processing in GC is a function of embeddedness in the amygdala-cortical loop

While our analyses do not suggest that Output and Input neurons are separate neuron types, our ultimate hypothesis was that the “embeddedness” of GC neurons within the reciprocally-connected network—whether they influence BLA (Output neurons), are influenced by BLA (Input neurons), both, or neither—determines their involvement in taste processing. Having conservatively labeled each neuron’s functional role, we tested this possibility; specifically, we tested whether we can discern a relationship between pGLM label and the magnitude of the stimulus-specificity (i.e., more distinctive responsiveness to different tastes) and palatability-relatedness (i.e., correlation with the hedonic value of the tastants) of the neuron’s taste response—the basic variables that describe involvement in taste processing (1–4,10,23–25,48,49).

Our analyses did in fact reveal significant differences in both taste-coding variables as a function of subpopulation (**Fig. 6A**): GC Input neurons (again, these are those shown by pGLM to be influenced by BLA, with no anatomical specifics implied) showed higher magnitudes of both taste specificity and palatability relatedness than did GC Output neurons. Despite the status of GC as a primary sensory cortex, the recipient of ascending taste input from thalamus (14,19,50,51), involvement in taste processing (as measured by taste specificity and palatability relatedness of taste responses) appears to depend on receiving “feedback” input from BLA.

**Fig. 6).**
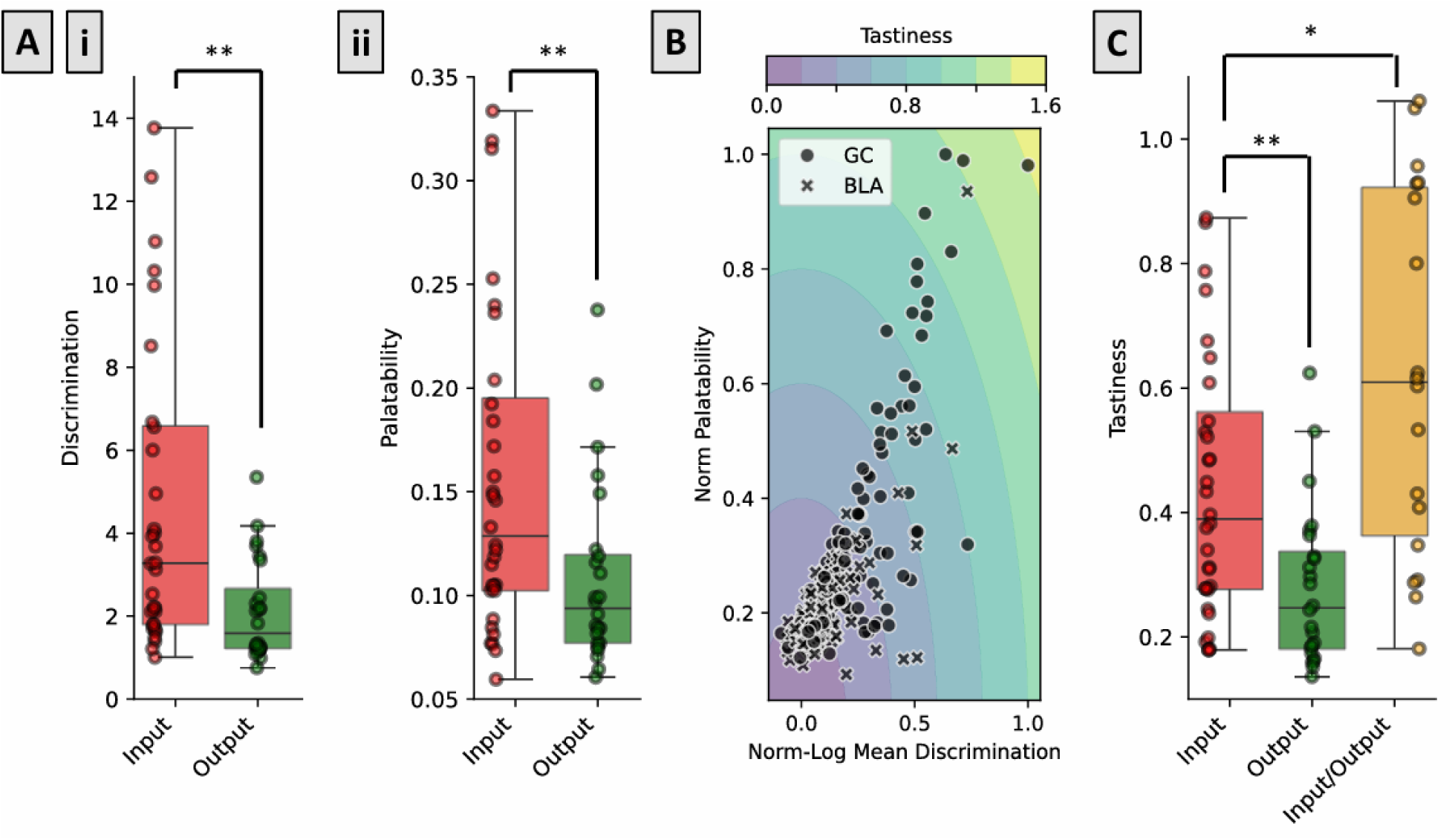
GC neurons that are embedded in the amygdala-cortical loop are the ones that most strongly process tastes. **A)** Both taste discriminability **(i)** and palatability-relatedness **(ii)** of GC Input neurons (those influenced by BLA activity), measured in the standard manner (see Methods), are significantly stronger than that in GC Output neurons. ** = p<0.005. **B)** Discriminability and Palatability are significantly correlated for GC neurons (Spearman’s Rho=0.73, p<0.001) and hence can be combined into a measure ( “Tastiness”) that represents a single measure of involvement in taste processing. **C)** Comparison of tastiness among subpopulations confirms the results of A (higher tastiness in Input neurons than Output neurons), but also shows that the small population of GC Input/Output neurons produces responses with significantly higher tastiness than even the Input population (2-sample Student’s T-test, ** = p<0.005, * = p<0.05)

As taste discriminability and palatability are (necessarily) non-independent, correlated variables (**Fig. 6B**, Spearman’s Rho=0.73, p<0.001), it was useful to aggregate and summarize the results shown in **Fig. 6A** into a single measure. We used for this summary the vector norm (the magnitude of an arrow drawn to a point in 2D-space, see Methods) of normalized discriminability and palatability coding, an aggregate measure of taste-processing involvement that we here refer to as “tastiness.” An analysis of neuron tastiness (**Fig. 6C)** confirms the findings of **Fig. 6A**: GC Input neurons show significantly higher tastiness than GC Output neurons (note that this difference was not an artifact of lower firing rates in the Output neurons—we observed no significant link between firing rates and the metrics tested above; Spearman’s Rho: Firing Rate vs. Discriminability = 0.1, Firing Rate vs. Palatability = 0.0).

But the most rigorous test of our central hypothesis (that involvement in taste processing depends upon being “embedded” in the amygdala-cortical loop) involves an analysis of the specific set of neurons that are most deeply so embedded—that is, the GC neurons that are categorized as both influencing and receiving influence from BLA (i.e., “Input/Output” neurons). If this hypothesis is correct, we should expect that these neurons will exhibit the strongest tastiness—even stronger than that of GC Input neurons. Indeed, this proved to be the case: the tastiness of GC Input/Output neurons is significantly higher than that of GC Input neurons (Student’s t-test p=0.01).

Note that the division of neurons into functional Input and Output types was made by a model that was blind to taste coding in those responses, and therefore our results provide a conservative, risky test of the importance of BLA feedback in taste responsiveness. Thus they confirm our prediction that the degree of “embedding” in the reciprocal circuit determines its degree of involvement in taste processing (**Fig. 6C**), a result that is particularly striking given the weak “tastiness” of GC Output neuron responses.

## Discussion

This study reveals GC taste processing to centrally involve dynamical activity of the amygdala-cortical loop. Following an initial transient in which they fire independently of one another, the flow of influence between BLA and GC becomes one of coupled, asymmetric “turn-taking.” Influence in the BLA→GC direction appears dominant during the middle epoch, when BLA taste responses are palatability-related and GC taste responses are not; GC→BLA influence, meanwhile, spikes immediately afterward, at the time that GC emits a signal reflecting the reaching of a consumption decision (almost certainly based on the prior input from BLA). At the single-neuron level, any individual GC neuron’s involvement in taste processing—measured in terms of the taste-specificity and palatability-relatedness of its taste responses—is shown to be a function of its embeddedness in this reciprocal circuit.

Thus, we conclude that the BLA-GC dyad, formed soon after taste delivery, processes tastes on the basis of direct recurrent connectivity between the two regions, collectively transitioning through quasi-stable states. The following sections discuss this proposal in further detail, specifically considering: 1) how stimulus-evoked metastable dynamics seen in the amygdala-cortical circuit are well-described as a distributed attractor network; 2) whether this conceptualization can be generalized to describe work from other sensory systems; and 3) the fact that the “flow of influence” in tightly connected dynamical systems is likely to be asymmetric, with distinct dynamics of influence acting in concert with one another, And 4) a brief consideration of output pathways and possible role of this loop in other proposed functions of GC.

### Taste processing evolves through the functioning of an amygdala-cortical joint attractor network

The metastable population dynamics that make up GC taste responses have already been successfully modeled as an attractor network (52–55). Our current results reveal this attractor network to be made up not just of GC, but of the dyad of GC and BLA. In such a joint attractor network, there is no “source” of transitions, which are triggered by coordination of the noise in the system. As predicted by such a model, all but the initial transitions (which are “externally” driven by the stimulus) in taste-evoked activity happen, on average, with lags that are distributed around zero, despite the fact that the regions are connected by relatively long axons—a result that is difficult to explain using hierarchical/feedforward models which, by construction, contain sources and sinks of influence (56–58). Thus, we argue that taste input causes the BLA-GC system to switch from resembling a feedforward system to resembling a recurrent attractor network for purposes of taste processing.

This characterization of BLA and GC as a joint network is unlikely to be an isolated case. Our previous work has shown that at the single-neuron level, neurons in both the lateral hypothalamus (48) and the parabrancial nucelus of the pons (59) also show states in the taste-evoked responses. Indeed, as noted below, such multi-state responses are ubiquitous, and hence, suggest that BLA and GC are likely undergoing these state-transitions as part of the larger taste circuit, especially given the dense recurrent long-range connectivity between most regions in the circuit (14,15). The fact that both GC and BLA are also emotional, interoceptive, and multisensory “hubs” (60,61) makes it reasonable to speculate that they couple with the olfactory circuitry, as well as frontal cortices, in appropriate functional contexts as well.

Given these considerations, we also expect that changes in internal state of the animal alters BLA-GC coupling. Indeed, GC taste responses change with the rat’s level of engagement in the task (31), a change accompanied by dramatic changes in the GC’s LFP profile suggesting a change in the functional connectivity of the larger network. It is likely that such a change may be driven by, or in concert with, the amygdalar (or frontal cortical) neurons, firing of which is sensitive to both internal state and task structures (62,63).

### Reciprocal interactions underlie processing in other sensory systems

Evidence supporting our proposal that reciprocal connectivity causes regions to “coalesce” into a single perceptual processing unit abounds in work on other sensory systems. Global Workplace Theory (GWT) (8,64), for instance, claims that sensory information is first transmitted in a feedforward manner to specific brain regions; activation of these regions then engages a distributed network via reciprocal projections, which essentially binds the regions together. Further, these different states reflect different stages of perceptual processing and transitions between stages are determined by different mechanisms. Importantly, their 2^nd^ state is driven by establishment of recurrent interactions, is internally driven (as opposed to being stimulus-bound), and reflects “conscious” stimulus processing. This mirrors taste research showing that processing in the 2^nd^ state in GC taste-evoked activity is used to make consumption decisions (3,55,65–68). And finally, that perturbation of activity in this 2^nd^ state most strongly affects stimulus-related decision making in both the GWT-related visual blocking task (8) and taste responses (25). Similar reports of circuits undergoing a state transition from an initial feedforward to a recurrent (core processing) state can be found for more visual processing studies (7,9), as well as somatosensory (69), auditory (70), and olfactory processing (71).

### The division of function(s) between BLA-GC directional influences

While the fact that BLA and GC progress through taste processing in lock-step is itself evidence of the importance of the reciprocal connectivity between them, more straightforward evidence is found in the asymmetries in how each region influences the other. It is actually not unusual (quite the opposite) for recurrently interacting systems to contain such asymmetric influences. Examples of dynamical systems in which feedforward and feedback interactions have different strengths include half-center oscillators (72), Lotka-Volterra (prey-predator) models (73), and reservoir networks (74). Furthermore, researchers have reported asymmetric interactions analogous to our own in data collected from visual system recordings (17,36,75).

The “turn-taking” dynamics of the amygdala-cortical interaction we have observed in this study accord well with classic stimulus-valuation studies, including those performed by Damasio (37) and Schoenbaum (38), which have been interpreted as suggesting that amygdala, after playing a primary role in determining the value of a stimulus, provides this information to cortex which then uses the information to generate a behavioral decision. Our GrCa analysis provides the first representation of this “signal” being emitted by GC, but it is worth noting that Forseth et al. (76) observed a similar transient “signal” emanating from somatomotor regions approximately 250ms prior to the onset of response articulation in the above-discussed visual-naming task.

In the reverse direction, it is tempting to speculate that the tonic GC→BLA influence in the γ frequency range powers the processing attributed to BLA, perhaps by sending taste quality information to BLA while BLA returns palatability information, with the two regions co-evolving in this manner to converge to a decision (77). While GC neurons that influence BLA activity without themselves being influenced by BLA activity (i.e., GC Output neurons) showed weak taste coding, the above conclusion can nonetheless be easily reconciled with previous work (20) showing that a subset of GC→BLA projection neurons have more discriminatory taste-responses compared to the average of non-projecting neurons, in response to aversive tastants vs palatable tastants. This matches well with our data if we remember that the bidirectional GC Input/Output neurons will only appear as GC→BLA projection neurons if bidirectional projecctions are not considered.

### Output pathways and other GC functions

In the context of previous work (3,4,10,25,37,38,78), the current findings can be interpreted to demonstrate that amygdala-cortical interactions culminate in the generation of consumption decisions. As to what happens next—how exactly the consumption decision is behaviorally enacted—our work is agnostic. Basic consumption-related behaviors are actuated by a central pattern generator in the brainstem, the Reticular Formation (RF; (79–81)), and thus it is reasonable to speculate that information flow from GC to RF is capable of driving these behaviors. A direct pathway connects GC and RF (82,83), but there are in addition multiple indirect feedback pathways linking the two structures (84,85). A recent proposal suggests that palatable and aversive taste behaviors may be instigated by separate feedback pathways leading through different amygdalar structures (86,87). While the connection between this work and ours is currently unclear (the proposal of separate behavioral pathways relies on the suggestion that spatially separated populations of GC neurons code the different tastes—a finding that most labs do not observe; (1,68,88–91), future work will illuminate that connection.

Another subject of future work is the involvement of the amygdala-cortical loop in other GC functions. GC responses have been shown in multiple contexts to represent post-ingestive factors; relatedly, recent work has highlighted the role of GC in interoceptive processes (60,92,93). While for the current study we expressly used an experimental protocol that minimized the influence of these variables (delivering small aliquots of fluid such that rats neither satiated nor ingested substantial numbers of calories, and randomizing presentation order and timing such that rats could not anticipate any delivery) such that taste responses were stable across entire sessions (94), it is entirely likely that the kind of amygdala-cortical dynamics exposed here are differentially invoked when a rat is adjusting its consumption on the basis of post-ingestive need. Even more generally, while the BLA-GC connection is likely stable during our roughly hour-long recording sessions, it will inevitably be modulated by long-term brain-states given that neural networks display multi-scale, context-dependent dynamics and interactions (95–97), making this an exciting avenue for future research.

## Conclusion

This study furthers our understanding of the role of inter-region interactions in sensory processing by showing that cortico-amygdalar interactions subserve taste processing and decision-making. We speculate that these dynamics are potentially a function of the larger taste circuit including the brainstem (59). This taste network is initialized in a feedforward state but coalesces into a reciprocally interacting network that evolves toward a point shown previously to generate the behavioral output (3,25). Our investigation of the makeup of GC Input and Output neurons suggests that network interactions are strong determinants of neural encoding and response properties. All results presented herein align with multiple previous studies across sensory systems as well as investigations of GC-BLA interactions during learning, allowing us to conceptualize these previous results in a unified theoretical framework.

## Methods

### Experimental Design and Statistical Analyses

#### Subjects

Adult, female Long-Evans rats (n = 6; 300–350g at time of electrode implantation, Charles River Laboratories) served as subjects in our study (we have observed no sex differences in the basic cortical dynamics of taste responses between male and female rats, and therefore use female Long-Evans rats because they are, in our hands, calmer than males). The rats were housed in individual cages in a temperature-and humidity-controlled environment under a 12:12 hr light:dark cycle, given ad libitum access to food and water prior to the start of experimentation, and weighed daily following surgery to ensure that they never dropped below 80% of their pre-surgery weight. All experimental methods were in compliance with National Institutes of Health guidelines and were approved in advance by the Brandeis University Institutional Animal Care and Use Committee.

#### Electrode and intra-oral cannula construction

Custom microwire bundle drives, optimized for chronic (i.e., multi-day and multi-week) recording quality, were made with either 16 or 32 electrodes per recording site (design and construction details available at https://katzlab.squarespace.com/technology). Intra-oral cannulae—flexible tubing with a flanged tip and washer to ensure stability, connected to a plastic top complete with a locking mechanism—were built to allow the delivery of tastants directly onto the tongue (94).

#### Surgery

Rats were anaesthetized with an intraperitoneal injection of ketamine/xylazine cocktail (100mg/kg and 5.2 mg/kg respectively) and mounted in a stereotaxic instrument (David Kopf Instruments; Tujunga, CA) with blunt (atraumatic) ear bars. A midline incision exposed the skull and trephine holes (∼2 mm diameter) were drilled above BLA and GC. Microwire bundles were implanted 0.5mm above GC (coordinates: AP +1.4 mm, ML-5.0 mm, DV −4.4mm from dura) and BLA (coordinates: AP - 3.0mm, ML-5.0mm, DV-6.8mm from dura). Once in place, electrode bundles were cemented to the skull. Once electrode bundles were secured, an intra-oral cannula (IOC) was threaded through the masseter muscle (inside the zygomatic arch) to the space between the lip and gums, and the top of the cannula was cemented to the rest of the assembly with dental acrylic (94). The rat’s body temperature was monitored and maintained at ∼37°C by a heating pad throughout the duration of the surgery.

#### Acquisition of electrophysiological data

Electrophysiological signals from the micro-electrodes were sampled at 30 kHz using 32-channel analog-to-digital converter chips (RHD2132) from Intan Technologies, digitized online at the head stage and sampled jointly, along with signals from actuators marking tastant delivery, using an Intan RHD USB interface board (Part #C3100), which routed records to the hard drive of a PC for saving.

The experimental chamber was ensconced in a Faraday cage that shielded recordings from external electromagnetic influences.

#### Habituation and passive taste administration

Following their recovery from surgery, we habituated rats to the experimental chamber for 2 days, to the IOC/electrode harness for the next 2 days, and to passive water deliveries for the following 2 days, before beginning data collection. Starting with the second habituation day, we also placed rats on a mild water restriction schedule—20mL of water (not including the ∼4mL delivered during habituation sessions themselves) per day. This water restriction schedule was maintained till the end of the experiment. For the 2 final habituation sessions, we attached the rats to the taste delivery apparatus, and infused 120 pulses of distilled water (∼30μL per pulse; 20s inter-pulse interval) into the animal’s oral cavity through the IOC, and drove electrode bundles deeper (by 250 μm) into target structures.

By the end of this procedure, the tips of the electrodes lay within GC and BLA. We then recorded taste responses during 3-4 days of taste delivery sessions, between each of which the microwire bundle was driven down approximately 60 μm. During these sessions, Sucrose (0.3M), Sodium Chloride (0.1M), Citric Acid (0.1M), and Quinine (1mM), dissolved in ultra-pure water (∼30μL per pulse; 20s inter-pulse interval, 30 trials/tastant; ∼3.6mL total volume delivered) were delivered to passive rats (i.e., no behavior was required to elicit delivery). These concentrations were chosen to represent a range of hedonic values, and because they are known to evoke robust responses in both GC and BLA (2,24). We note that this volume has been shown to evoke stable neural responses in roughly hour-long sessions, arguing against evocation of post-ingestive effects during the recording (31).

#### Histology

In preparation for histology, rats were deeply anesthetized with an overdose of the ketamine/xylazine mixture. We perfused the rats through the heart with 0.9% saline followed by 10% formalin and harvested the brain. The brain tissue was incubated in a fixing mixture of 30% sucrose and 10% formalin for 7 days before being sectioned into 50μm coronal slices on a sliding microtome (Leica SM2010R, Leica Microsystems). Sections containing the electrode implant sites around GC and BLA were imaged at 2x.

## Data and statistical analyses

The analysis of data and statistical tests were performed using custom written software in Python as described below.

### Local Field Potential processing and analysis

#### Filtering / power and phase extractions

LFPs were extracted from broadband digitized signals using a 2nd order bandpass Butterworth filter (1-300Hz), to de-emphasize spiking and emphasize frequencies typically of interest in such data. In order to avoid contamination from noise/artifacts on noisy/broken channels, only channels containing isolable single neurons (see below) were used for analyses. Signal was down sampled to 1000Hz to reduce computational load for downstream analyses.

We note that this filtering process is sufficient for analysis up to the maximum frequency shown in the Results section (that is, 100Hz), and is unlikely to be the reason we don’t observe effects above 60Hz. First, a maximum frequency of 300Hz is sufficient for a Nyquist frequency for accurately detecting signals up to 150Hz. Secondly, we use a Butterworth filter, which is “maximally flat” (see scipy.signal.butter, Virtanen et al., 2020) within the pass band and is even permissive of frequencies higher than 300Hz. Hence, signals with frequencies 100Hz or lower are unlikely to be lost due to filtering.

#### Spectral Granger Causality

Spectral Granger Causality was assessed between pairs of electrodes in GC and BLA using the Spectral Connectivity package (30). We selected for this analysis the electrodes from which activity was most similar to the mean activity for the region (smallest mean-squared error relative to the mean phase across all channels for each region; this selection of channels was constant for all trials in a single analyzed session, and the same set of channels was used for all frequencies), to ensure reliable representation of each.

To satisfy the normality assumption for use of Granger Causality, signals from both electrodes were first preprocessed according to previously established criteria to remove outliers and “detrend” the data (99). Briefly:

1. Any trials with data points exceeding 3 Median Absolute Deviations calculated using all the trials, were removed
2. Linear detrending was performed on a single-trial basis (scipy.signal.detrend)
3. The temporal (single-trial) mean was subtracted from the data, and all data divided by the temporal standard deviation
4. Trial-averaged mean was subtracted, and all data divided by the trial-averaged standard deviation

Following preprocessing, the Augmented Dickey-Fuller test was run on each channel to confirm stationarity (normality) of the signal. All preprocessed data passed the test and hence was included in further analyses.

Using the preprocessed data, spectral granger causality was calculated using multitaper methods implemented in the Spectral Connectivity package. The default set of tapers (Slepian) was used. Specific parameters included:

- “sampling_frequency” = 1000 (Hz),
- “time_halfbandwitch_product” = 1
- “time_window_duration” = 0.3 (sec)
- “time_window_step” = 0.05 (sec)

Significance of the granger causality for each frequency-time bin on a single-session basis was obtained using a shuffling procedure. Granger Causality was recalculated for 500 sets of data where trial labels were shuffled (mismatched) between BLA and GC to generate a null distribution of “non-specific” Granger Causality between the two regions. Deviation of the value in each frequency-time bin from its respective null distribution was taken to indicate a significant interaction in that bin. Multiple comparisons were addressed using the Bonferroni correction with a base alpha of 0.05 with each frequency-time bin treated as a comparison. We further supplemented this “bin-level” testing as below.

#### Cluster Permutation Testing

To perform statistical testing on the Granger Causality results while minimizing issues of multiple comparisons, we opted to use a cluster-based permutation test (100,101) to detect the largest statistically-significant deviation in the time-series for the 2-12Hz and 20-60Hz frequency bands. Briefly, the procedure involves:

1. Z-scoring the time-series on which testing needs to be done (in our case, this is the mean significance of Granger Causality for a frequency band)
2. Assessing the summed value of the largest cluster (continuous chunk) above a specific threshold for the absolute of the z-scored time-series (we used a threshold of Z = 2). This value is used as the test statistic.
3. This summed value of the largest cluster is then recalculated from shuffled data to obtain a null-distribution for the test statistic.
4. If the actual value of the statistic significantly deviates from the null-distribution, the null hypothesis is rejected (alpha=0.05).

#### Granger Causality Similarity Analysis

To analyze similarities in the dynamics of the directional Granger Causality timeseries, we performed Principal Component Analysis on the Granger Causality significance timeseries for 0-100Hz, reducing each multivariate timeseries down to 3 components to mitigate overfitting in the following regression analyses. All comparisons were then performed on these dimensionally reduced timeseries. We next performed 1) Linear Regression between the two timeseries, 2) Linear regression between circularly (temporally) shuffled versions of the two timeseries (as a negative control / context for poor prediction), 3) Linear Autoregression, and 4) non-Linear Autoregression (Multilayered perceptron regressor, (102). For both forms of autoregression, we used the previous 3 timepoints to predict the current one. To obtain an estimate of the uncertainty of prediction, we performed bootstrapping of the data with 500 reruns of model fitting and prediction.

## Analysis of Spiking Activity

### Single Unit Isolation

Spikes from electrophysiological recordings were sorted and analyzed off-line using an in-house Python-based pipeline (103). Putative single-neuron waveforms with >5:1 signal-to-noise ratio (median absolute deviation) were sorted using a semi-supervised algorithm: recorded voltage data were filtered between 300-3000Hz, grouped into potential clusters by a Gaussian Mixture Models (GMM) fit to multiple waveform features; clusters were then labeled and/or refined manually (to increase conservatism) by the experimenters.

### Single Unit Evoked Response Characterization

#### Evaluating single-neuron response taste specificity

To statistically determine the degree to which a single neuron’s response contained taste-specific information, firing rates were estimated by convolving spike-trains with a rectangular window (length: 200ms). A one-way ANOVA (between tastes, across trials) was then run on each window to identify if the response of a single neuron to one taste was different from its responses to any other tastes at that timepoint. The (taste) specificity of a single neuron was defined as the average of the ANOVA effect sizes for 2000ms of post-stimulus activity.

#### Evaluating single-neuron response palatability-relatedness

To statistically determine the degree to which a single neuron’s response reflected the hedonic value of the stimuli delivered, we estimated firing rates as described above, and calculated the Pearson’s correlation coefficient between the evoked firing rates and the palatability ranks of the tastants at each timepoint. Palatability ranks — sucrose (1) > NaCl (2) > citric acid (3) > quinine (4) — directly reflected consumption of (earlier-run squads of) rats in a Brief Access Task (see (2)). This ordering is canonical, and has been replicated in many studies and with multiple measures of stimulus appreciation (94,104,105). The palatability of a single neuron was defined as the average of the absolute correlation value (as the correlation can also be strongly negative) for 2000ms of post-stimulus activity.

### Calculating “Tastiness”

To combine taste specificity and palatability into a single metric, we first log-transformed specificity to make the scales of specific and palatability more similar. This is because palatability as an absolute value of a correlation is bounded between [0,1] whereas specificity as an effect size has the range [0, ∞). Log-Specificity and Palatability were then Min-Max normalized following which the L2-norm of the vector generated by the normalized metrics (normalized log-specificity, normalized palatability) was calculated on a single-neuron basis and defined as “tastiness”.

#### Determination of putative pyramidal vs inter-neurons

From the average of filtered spikes for each putative single-unit, we determined the time taken from the trough to the next peak. As established previously, inter-neurons and pyramidal neurons can be reliably separated using this “rise time” (4,45–47). The specific threshold varies depending on the filtering frequencies used. We used a value of 0.45ms based on previous work using the same bandpass filter frequencies as we have used here (4,47)

### Changepoint Modeling of Population Activity

#### Model Fitting

Changepoint modelling of neural activity was performed as described previously (106) using the pytau library (https://github.com/abuzarmahmood/pytau) written in Python. Briefly, spike-trains were binned into 50ms bins. Changepoint models with independent Poisson emissions were fit to these timeseries of binned spike-counts for BLA and GC populations separately using automatic differentiation variational inference (ADVI) functionalities implemented in the pymc3 probabilistic programming library (107). Once fit, the procedure returns: 1) the estimate of the posterior distribution for single-trial population changepoint locations, and 2) the emission (firing) rates for each neuron during every state. Since ADVI utilizes spherical Gaussians as the surrogate distribution, we use the mode of the distribution for individual parameters as the “best estimate” of the parameter for downstream analyses. Models with 4 states were fit to 2000ms of post-stimulus activity. Please refer to (106) for more details on the process of model/state-number selection.

#### Calculation of Transition Lags

Lags between BLA and GC transitions were calculated on a single-trial basis by subtracting the “best estimate” (see above) of the matched transition (that is, transition 1 in GC was compared to transition 1 in BLA for any given trial) between models fit to each region. This resulted in a distribution of lags for each recording session (120 values, 1 for each trial in a session). The means of each session’s distribution were used to generate a distribution of average lags for each transition across all the recording sessions. Finally, a grand average of each transition’s distribution of average lags was calculated. The significance of the deviation of this grand-average lag from zero was calculated by bootstrapping at the level of trials for each session with a total of 5000 permutations. The 95% confidence interval of each grand average was calculated using these bootstrapped distributions, and deviation of zero from this 95% interval was taken as a significant difference.

### Generalized Linear Modelling

As has been shown previously, Granger Causality methods perform poorly on spike-train data due to the strong violation of the stationarity assumption used by the autoregressive processes on which Granger Causality is based (108,109). This issue is addressed by General<u>ized</u> Linear Models (GLM; distinct from <u>General</u> Linear Models which also follow the stationarity/normality assumption) which can model Poisson-distributed data, and furthermore can model temporal dependencies explicitly over a large range of timescales (40,57,110,111).

#### Model Specification and Model Fitting

The GLM included 3 sets of filters for each neuron modelled (see **Fig. 4A** for schematic):

- <u>Spike History Filter</u>: Captures how a neuron’s past spiking activity influences its current firing rate. This can model refractoriness (reduced probability of firing immediately after a spike) and bursting behavior.
- <u>Stimulus Filter</u>: Models how external stimuli drive neural activity.
- <u>Coupling Filter</u>: Captures how the activity of other neurons influences the target neuron, modeling functional connectivity.

The duration of the spike history and coupling filters was set to 100ms, while the duration of the stimulus filter was set to 300ms, based on previous modelling work (40). To reduce number of fit parameters and mitigate over-fitting of the models, raised cosine-basis functions were used to parameterize the filters (40,41,110): a basis of 10 functions was used for every filter, with peaks distributed logarithmically to model fast and slow timescales with appropriate temporal resolutions.

The conditional intensity of the neural activity was hence defined as:

λ(t) = exp(β_0_ + β_hist * h(t) + β_stim * s(t) + β_coup * c(t))

- λ(t) is the firing rate at time t
- β₀ is the baseline firing rate
- h(t) represents the neuron’s own spike history
- s(t) represents the stimulus
- c(t) represents the coupling with other neurons
- β terms are the corresponding weights

The Python “statsmodels” library was used for model fitting (112).

Quality of fit was assessed in two ways. First, by comparing the log-likelihood of models fit to actual data with models fit to data where the trial labels were shuffled for neurons in the ensemble being fit, which would only allow the model to learn an “average” coupling filter between the neurons in the model being fit. Second, we compared Spearman’s Rho between predicted peri-stimulus time histograms (PSTH) of a given neuron with PSTHs predicted for 1) the neuron the model was fit to (actual data), 2) a different neuron (Unit Shuffled), and 3) temporally scrambled data for the same neuron (Circular Shuffle).

Once models were fit, putative inter-region connections were determined using the coupling filters. The fitting process returns p-values for each coefficient in the coupling filter (which, in our case due to the parametrization of the filter using basis functions corresponded to the scaling factor of each function in the basis). We asserted that a coupling filter indicated a statistically significant interaction if a p-value in the filter was below alpha=0.05 following a Bonferroni correction (corrected p-value = 0.05 / 10 = 0.005).

#### Assessment of overlap between input and output populations

We assess whether the overlap observed between subpopulations labelled as “Input” and “Output” in GC and BLA was different from that expected by chance using Monte Carlo simulations. Essentially, keeping the number of Input, Output, and Total number of neurons in each region constant, we label neurons as Input or Output randomly and assess the number of neurons with both labels. We perform 10,000 simulations for each region to generate a null distribution for this overlap as expected by random assignment. Deviation of the observed value of overlap from the 95% confidence interval of this distribution would indicate that the observed value is significantly different from that expected by chance.

## Data Availability

We have structured our electrophysiology datasets in a hierarchical data format (HDF5) and are hosting the files on a university-wide network share managed by Library and Technology Services (LTS) at Brandeis University. These HDF5 files contain our electrophysiology recordings, sorted spikes, single-neuron and population-level analyses (and associated plots and results). These files are prohibitively large to be hosted on a general-purpose fileshare platform - we request anyone interested in our datasets to contact the corresponding author, Donald Katz who can put them in touch with LTS in order to create a guest account at Brandeis through which they can securely access our datasets (hosted on the katz-lab share at files.brandeis.edu).

